# Selection of representative genomes for 24,706 bacterial and archaeal species clusters provide a complete genome-based taxonomy

**DOI:** 10.1101/771964

**Authors:** Donovan H. Parks, Maria Chuvochina, Pierre-Alain Chaumeil, Christian Rinke, Aaron J. Mussig, Philip Hugenholtz

## Abstract

We recently introduced the Genome Taxonomy Database (GTDB), a phylogenetically consistent, genome-based taxonomy providing rank normalized classifications for nearly 150,000 genomes from domain to genus. However, nearly 40% of the genomes used to infer the GTDB reference tree lack a species name, reflecting the large number of genomes in public repositories without complete taxonomic assignments. Here we address this limitation by proposing 24,706 species clusters which encompass all publicly available bacterial and archaeal genomes when using commonly accepted average nucleotide identity (ANI) criteria for circumscribing species. In contrast to previous ANI studies, we selected a single representative genome to serve as the nomenclatural type for circumscribing each species with type strains used where available. We complemented the 8,792 species clusters with validly or effectively published names with 15,914 *de novo* species clusters in order to assign placeholder names to the growing number of genomes from uncultivated species. This provides the first complete domain to species taxonomic framework which will improve communication of scientific results.

## Introduction

Sequencing and computational advances have enabled the rapid, cost-effective recovery of cultivated and uncultivated bacterial and archaeal genomes. This has resulted in large-scale initiatives such as the Genomic Encyclopedia of Bacteria and Archaea (GEBA) producing thousands of isolate genomes assembled from type strains (Kyrpides et al., 2014; Mukherjee et al., 2017) and numerous studies reporting thousands of metagenome-assembled genomes (MAGs) recovered from a diverse range of environments (Parks et al., 2017; Chen et al., 2019; Pasolli et al., 2019). The availability of a large number of genome assemblies provides a wealth of genomic information for taxonomic classification (Konstantinidis and Tiedje, 2005a; Thompson et al., 2015; Garrity, 2016; Hugenholtz et al., 2016) which has been leveraged to reclassify specific bacterial lineages including the Epsilonproteobacteria (Waite et al., 2017), Actinobacteria (Nouioui et al., 2018), and *Lactobacillus* (Wittouck et al., 2019). We recently made use of this expansion in public genomes to produce the Genome Taxonomy Database (GTDB), a comprehensive genome-based taxonomy with bacterial and archaeal taxa circumscribed on the basis of monophyly and relative evolutionary divergence (Parks et al., 2018). Here we expand on this prior work by organizing all genomes encompassed by the GTDB into quantitatively defined species clusters based on the average nucleotide identity (ANI) to selected species representative genomes.

Many species definitions have been proposed for Bacteria and Archaea that consider different biological, ecological, and genomic aspects of these organisms (Cohan, 2002; Konstantinidis et al., 2006; Fraser et al., 2009; Bobay and Ochman, 2017). Here we were interested in an operational species definition that facilitates the automated assignment of genomes to species and that scales to large datasets in order to allow all available and forthcoming genomes to be organized into species clusters. This can be achieved using whole-genome ANI which has emerged as a robust and widely accepted method for circumscribing species (Ciufo et al., 2018; Chun et al., 2018), with 95% ANI found to recapitulate the majority of existing species (Konstantinidis and Tiedje, 2005b; Jain et al., 2018; Olm et al., 2019). Further supporting the use of ANI is its strong correlation with DNA-DNA hybridization which was considered the gold standard for circumscribing bacterial and archaeal species for decades (Goris et al., 2007). ANI is determined from the similarity of orthologous regions shared between two genomes (Konstantinidis and Tiedje, 2005a) and a number of methods have been proposed for calculating this statistic (Richter M and Rosselló-Móra, 2009; Varghese et al., 2015; Yoon et al., 2017). Here we made use of two recent advances in calculating ANI that allow tens of thousands of genomes to be organized into species clusters; a fast heuristic for approximating ANI (Mash: Ondov et al., 2016) and a computationally efficient approach highly correlated with the results of traditional methods (FastANI: Jain et al., 2018).

Species clusters can be formed in a number of ways based on the ANI between genomes. A common approach is to represent genomes as nodes in a graph with edges between genome pairs having an ANI ≥95%. A graph-based clustering method can then be applied to divide the graph into putative species clusters without *a priori* consideration of existing nomenclature or taxonomy (Verghese el at., 2015, Ondov et al., 2016, Rodriguez-R et al., 2018). By contrast, we have taken an approach that explicitly accounts for validly or effectively published species names, directly ties species clusters to nomenclatural type material where possible, and results in a single representative for each species cluster. Specifically, we identified genomes assembled from the type strain of the species and used these as the representatives of species clusters circumscribed using ANI. We believe the use of genomes assembled from the type strain is a pragmatic choice for circumscribing species given systematic efforts to sequence such strains (Kyrpides et al., 2014; Mukherjee et al., 2017) and the nomenclatural and taxonomic importance of type material (Parker et al., 2019). Genomes that were not assigned to a named species cluster were organized into *de novo* species clusters with representative genomes selected based on genome quality and acting as nomenclatural type material. This follows the recent proposal that gene sequences are suitable type material for Bacteria and Archaea to the extent that they allow for the unambiguous circumscription of taxa (Whitman, 2016; Konstantinidis et al., 2017; Chuvochina et al., 2019). The proposed species clusters encompass all public genomes within the NCBI Assembly database and have been incorporated into the Genome Taxonomy Database to provide a taxonomic framework where genomes have assignments at all ranks from domain to species (Parks et al., 2018).

## Results

### Identification of genomes assembled from type material

The proposed species clusters were determined from a dataset comprising 153,849 genomes obtained from the NCBI Assembly database (Kitts et al., 2016) and 153 previously recovered archaeal MAGs (Parks et al., 2018; **Fig. 1A**). Genomes assembled from type material were identified by cross referencing strain identifiers at NCBI with the co-identical strain identifiers at LPSN (List of Prokaryotic Names with Standing in Nomenclature; Parte 2018), BacDive (Reimer et al., 2019), and StrainInfo (Verslyppe et al., 2014). Unfortunately, associating genome assemblies with nomenclatural types remains a challenge with currently available resources (Federhen, 2015). Type material is referenced by the unique strain identifiers at each culture collection resulting in a list of co-identical strain identifiers (e.g., *E. coli* ATCC 11775 = CCUG 24 = … = NCTC 9001). Nomenclatural resources such as LPSN and BacDive have no easy mechanism for maintaining a complete list of these co-identical identifiers and the strain identifiers associated with genomes at NCBI are largely the responsibility of individual submitters. Consequently, genomes may be identified as being assembled from type material at only a subset of nomenclatural resources and do not always agree with the nomenclatural status of genomes at NCBI (**Fig. 1B**). This latter situation is conflated by genomes at NCBI being annotated as assembled from type material if a genome has been ‘effectively published’ (e.g. *Clostridium autoethanogenum* DSM 10061), but the species name not formally validated (Parker et al., 2019). We considered a genome to be assembled from type material if any of its strain identifiers at NCBI could be matched with a co-identical strain identifier at LPSN, BacDive, or StrainInfo. This results in 8,665 genomes spanning 7,104 species being identifies as assemblies from the type strain of a species (**Supp. Table 1**).

**Figure 1.**
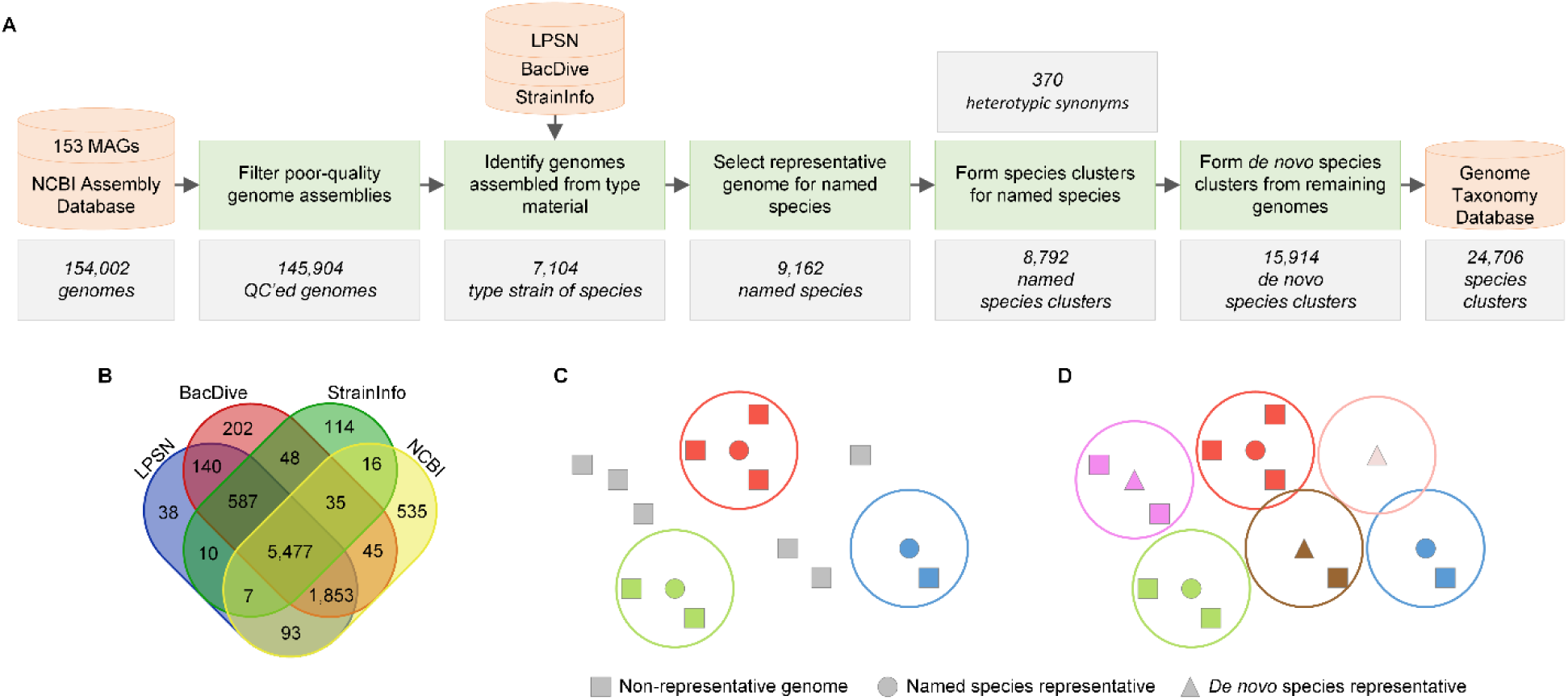
Overview of workflow for organizing genome assemblies into species clusters. (**A**) A dataset of 152,288 bacterial and 2,661 archaeal genomes was obtained from the NCBI Assembly database and supplemented with 153 archaeal MAGs. These genomes were filtered to remove 8,098 low-quality genomes. LPSN, BacDive, and StrainInfo were cross referenced with the species and strain information at NCBI to identify genomes assembled from the type strain of a species. A single representative genome was selected for each of the 9,162 validly or effectively published species names associated with one or more genomes in the dataset, with preference given to genomes assembled from the type strain of the species. Clusters were formed for named species based on the ANI between species representatives and all other genomes. This resulted in 8,792 species clusters due to the formation of 370 synonyms between closely related species representatives. Genomes not assigned to a named species cluster were formed into 15,914 *de novo* clusters using a greedy clustering algorithm which prioritized high-quality genome assemblies. The resulting 24,706 species clusters encompass all 145,904 quality-controlled genomes and have been incorporated in the taxonomy of GTDB release R04-RS89. (**B**) Overlap between genomes determined to be assembled from the type strain of the species at LPSN, BacDive, StrainInfo, and NCBI, highlighting the incomplete nature of co-identical strain identifiers between these different nomenclatural resources. (**C**) Conceptual illustration of genomes with the ANI between genomes depicted by their Euclidean distance. Three selected representative genomes for validly or effectively published species names are shown as circles with their circumscription radii depicted by larger circles. Genomes assigned to each named species representative are shown by squares of the same color. Genomes not assigned to a representative are shown in grey. (**D**) As per panel C with the additional selection of three *de novo* representative genomes which result in all genomes being assigned to a species cluster.

### Representative genomes for named species

Species clusters were formed by selecting a single representative genome for each of the 9,162 validly or effectively published species names associated with one or more of the 145,904 quality-controlled genomes (**Supp. Fig. 1**; **Fig. 1A**). Of these species, 5,942 (64.9%) consisted of a single genome which was selected as the species representative. The remaining 3,220 species were comprised of multiple genomes and the representative was selected by giving preference to genomes i) assembled from the type strain of the species (2,632 species), ii) annotated as being assembled from type material at NCBI (123 species), iii) designated as a reference or representative genome at NCBI (220 species), or iv) assembled from the type strain of a subspecies (8 species). In 1,506 cases, multiple potential representative genomes were still available within a genome category (i.e., multiple genomes assembled from the type strain of the species) and the representative was selected by considering NCBI metadata, the ANI between genomes, and in a small number of cases by manual investigation (219 species; see Methods). Overall, 7,104 of the 9,162 (77.5%) species were represented by a genome assembled from the type strain of the species (**Fig. 2A**; **Supp. Table 2**) demonstrating the success of initiatives such as GEBA that aiming to sequence all available type strains (Kyrpides et al., 2014).

**Figure 2.**
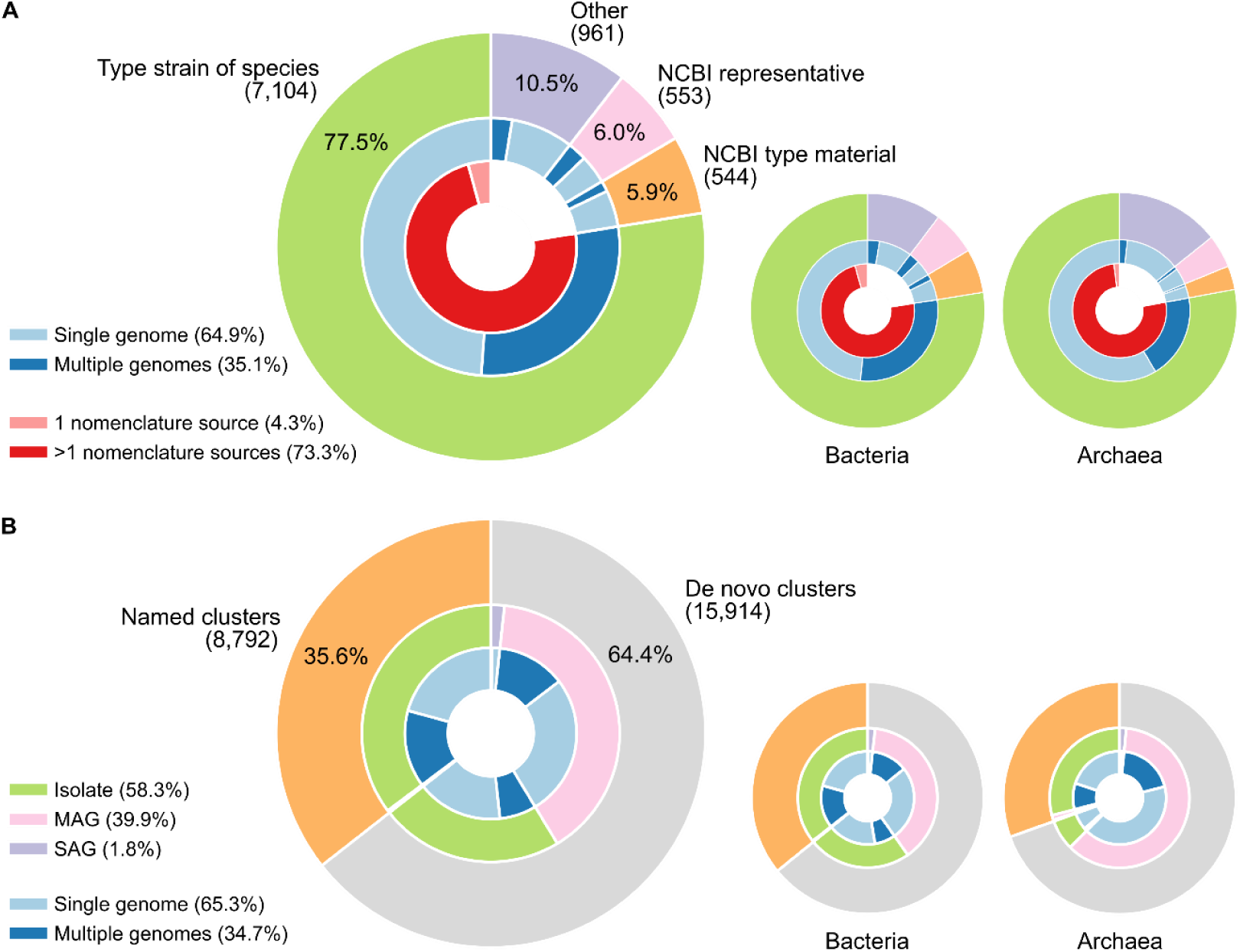
Properties of genomes selected as species representatives. (A) Representative genomes selected for the 9,162 validly or effectively published species names. Outer ring indicates the proportion of genomes selected from different metadata categories. Middle ring indicates the proportion of species within each metadata category that consist of either a single genome or multiple genomes. Inner ring indicates the proportion of genomes designated as being assembled from the type strain of species based on co-identical strain information from either single or multiple nomenclature resources (i.e. LPSN, BacDive, StrainInfo). (B) Representative genomes for the 24,706 species clusters circumscribing the 145,904 quality-controlled genomes. Outer ring indicates the proportion of named and *de novo* species clusters. Middle ring indicates the proportion of species with an isolate, MAG, or SAG as the representative genome. Inner ring indicates the proportion of species clusters consisting of either a single genome or multiple genomes. Inset charts show results for bacterial and archaeal species using the same color scheme and layout.

### Circumscribing named species clusters

Species clusters are circumscribed based on the ANI and alignment fraction (AF; i.e. percentage of orthologous genes) between genomes. The ANI circumscription radius for each named species representative was set to 95% ANI except if two representatives had an ANI >95% (**Fig. 3A** and **3B**). In such cases, the ANI radius of a representative was set to the value of the closest representative up to a maximum of 97%, with species representatives having an ANI >97% considered synonyms (**Fig. 3C**; *see below*).

**Figure 3.**
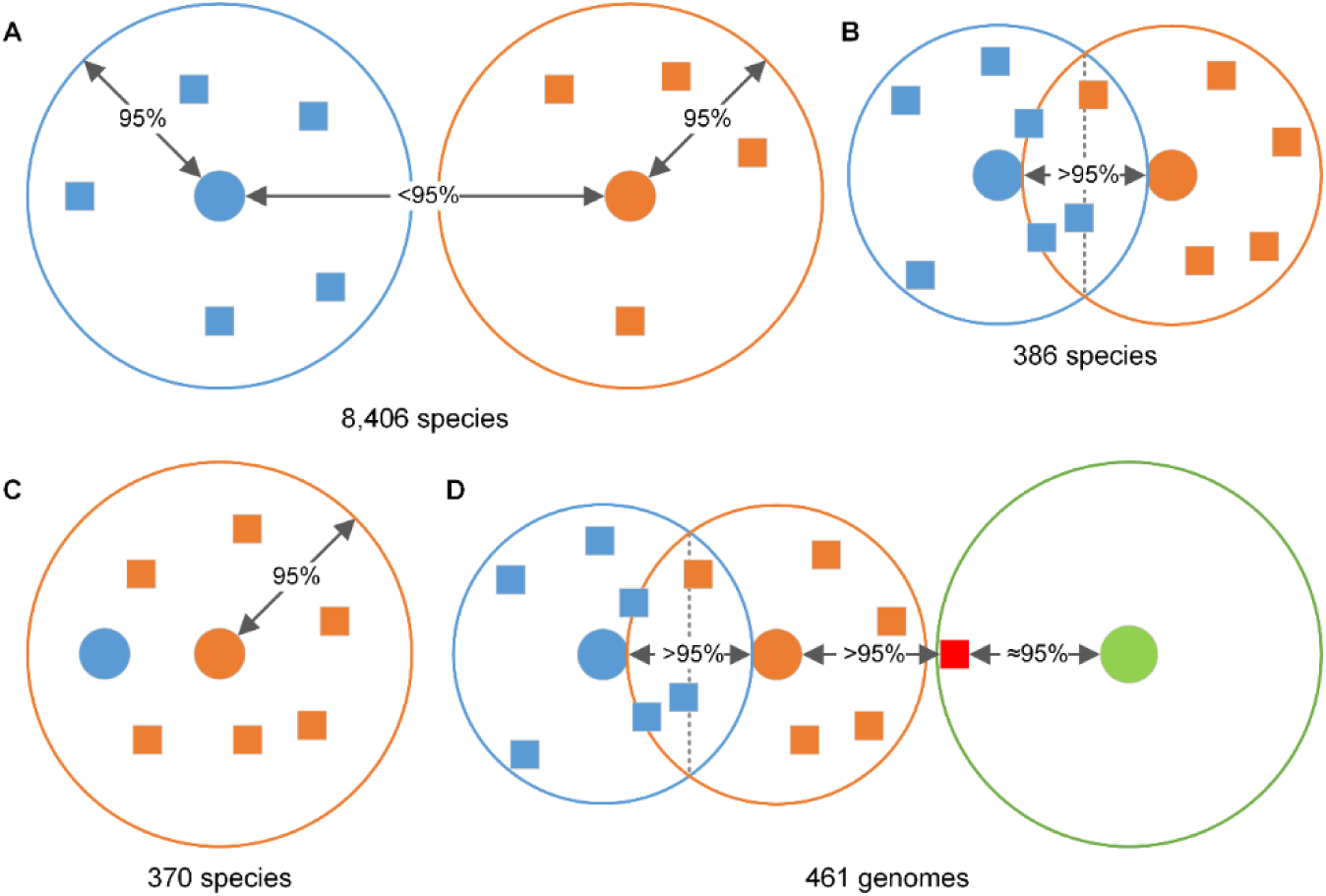
Illustrative examples of circumscribing species for varying ANI values between species representatives. The ANI between genomes is depicted by their Euclidean distance. Representative genomes are shown as circles with their circumscription radius depicted by larger circles. Genomes assigned to each representative are shown by squares of the same color. (A) Representative genomes with <95% ANI have a circumscription radius of 95%. (B) Representative genomes with an ANI between 95% and 97% have a circumscription radius equal to the ANI between the representatives. (C) Representative genomes with an ANI >97% are considered synonyms and only the representative of the species with priority retained. (D) Transitive situation illustrating a genome (shown in red) that is not within the ANI circumscription radius of the closest representative (shown in orange) and is therefore not assigned to any representative even if it is within the circumscription radius of another representative (shown in green).

The resulting ANI circumscription radii were then used to form species clusters for the 8,792 non-synonymous representative genomes (**Fig. 1C**). For each of the 137,112 remaining genomes in the dataset, the closest representative with an AF >65% (Konstantinidis and Tiedje, 2005b; Goris et al., 2007; Varghese et al., 2015) was determined and the genome assigned to this representative if it was within its ANI circumscription radius. This resulted in 104,763 (77.8%) genomes being assigned to named species clusters, with the most abundant species reflecting highly sequenced human-associated microorganisms (**Supp. Table 3**). The majority (87.3%) of assigned genomes only satisfied the ANI circumscription radius and 65% AF criterion for a single species representative. Genomes meeting the assignment criteria of multiple representatives were primarily classified as *Escherichia flexneri* (68.1%), *Escherichia dysenteriae* (9.0%), or Neisseria meningitidis_B (8.0%) (**Supp. Table 4**). In a small number of instances (461 genomes, 0.44%), a transitive situation arose whereby a genome was not within the ANI radius of the closest representative while being in the ANI radius of one or more other species representatives (**Fig. 3D**). These genomes were left unassigned in order to reflect the reduced ANI radius of species in their local phylogenetic neighbourhoods, which were almost exclusively within the genera *Escherichia* (95.2%) and *Serratia* (4.3%).

### Synonyms in the Genome Taxonomy Database

The 370 species reclassified as synonyms because they have an ANI >97% to another species with naming priority represents a practical compromise between having a quantitative species definition and retaining the majority (8,792 of 9,162; 96%) of species with validly or effectively published names (**Supp. Tables 5** and **6**). The necessity for ANI-defined synonyms was greatest in taxa of medical importance such as *Brucella* (Ficht, 2010), which comprises nine species as defined under the NCBI taxonomy (*B.* melitensis, *B. vulpis, B. ovis, B. canis, B. neotomae, B. suis, B. ceti, B. abortus*, and *B. microti*). These are reclassified as a single species, *Brucella melitensis*, in GTDB as the ANI is >99.5% and AF >93% for the representatives of these synonymous species with the exception of the *B. vuplis* genome at 97.5% ANI and 90% AF (**Supp. Table 6**). The high genomic similarity of these genomes suggests that they should be classified as monospecific subspecies or biovars as previously proposed (Verger et al., 1985). Similarly, several other instances of proposed synonyms are supported by our ANI-defined approach and have been incorporated into the GTDB taxonomy (**Supp. Table 6**). These include *Mycobacterium africanum, M. bovis, M. caprae, M. canettii, M. microti, M. mungi, M. orygis*, and *M. pinnipedii* as synonyms of *M. tuberculosis* (Riojas et al., 2018); *Bacillus plakortidis* and *B. lehensis* as synonyms of *B. oshimensis* (Liu et al., 2019); *Burkholderia pseudomallei* as a synonym of *B. mallei* (Varghese et al., 2015), and *Halomonas sinaiensi* as a synonym of *H. caseinilytica* (Oren, 2017). Note that 192 (52%) of the 370 ANI-defined synonyms are not based on type strains because genome assemblies are not available (**Supp. Table 6**). The status of these synonymous species will need to be reassessed once type strain sequences become available.

### Establishing de novo species clusters

The 32,349 genomes not assigned to named species were organized into *de novo* clusters using a greedy clustering approach favoring the selection of high-quality genomes to represent each cluster (**Fig. 1D**). Selection of representative genomes consists of four steps: i) sorting genomes without a species assignment by estimated genome quality, ii) selecting the highest-quality genome as a representative of a new species cluster, iii) determining the species-specific ANI circumscription radius for the new species cluster, and iv) temporarily assigning genomes to the new cluster using the same ANI and AF criteria used for named species clusters. These steps were repeated until all genomes were assigned to a species cluster. Finally, non-representative genomes were re-clustered to ensure that they had been assigned to the closest *de novo* species representative. This resulted in 15,914 *de novo* species clusters with the majority being represented by a MAG (61.6%) and comprised of a single genome (68.8%; **Fig. 2B**; **Supp. Table 7**).

The majority (58.4%) of *de novo* representatives meet the high-quality completeness (≥90%) and contamination (≤5%) MIMAG criteria (Bowers et al., 2017) suggested as suitable for defining sequence-based type material (Whitman, 2016; Chuvochina et al., 2019). The number of representatives suitable for use as type material increases to 73.5% under the more lenient proposal of 80% completeness (Konstantinidis et al., 2017). However, if the presence of a near-complete 16S rRNA genes (≥1200 bp) is required as generally recommended (Konstantinidis et al., 2017; Chuvochina et al., 2019) the number of potential type material representatives decreases substantially (37.0% at 90% completeness; 39.8% at 80% completeness) owing to this multi-copy gene often being absent from MAGs (Parks et al., 2017). As expected, the 8,792 representatives for named species are generally of high quality with 96.4% being ≥90% completeness with ≤5% contamination and 85.1% also containing a near-complete 16S rRNA sequence.

### Landscape of proposed species clusters

Here we explore key properties of species clusters defined by the ANI to a representative genome. The majority of the 24,706 species clusters are comprised of a single genome (65.3%) and only 919 clusters (3.7%) consist of ≥10 genomes (**Fig. 4A**; **Supp. Table 3**). A substantial number of clusters are comprised exclusively of MAGs (39.8%; 9,839 species) with the majority of these being singletons (67.5%; 6,646 species). While ANI circumscription radii between 95% and 97% were used to retain published species names, 8,407 (95.6%) of the 8,792 named species clusters have a radius of 95% (**Fig. 3A**; **Supp. Table 8**). Notably, 97.3% of the 4,660 species clusters with ≥3 genomes form a clique at 95% ANI (i.e. all pairwise combinations of genomes have an ANI ≥95%) with all species forming a clique at an ANI of 93.5% (**Fig. 4B**; **Supp. Table 8**). Inter-species ANI values between representatives indicate that the majority of species within a genus are highly divergent from each other with >96.2% of the 927,064 comparisons having an ANI <90% (**Fig. 4C**). In contrast, the ANI to the closest intra-genus species for the 19,898 species representatives within genera with multiple species shows a fairly even distribution between 78 to 95% ANI (**Fig. 4D**).

**Figure 4.**
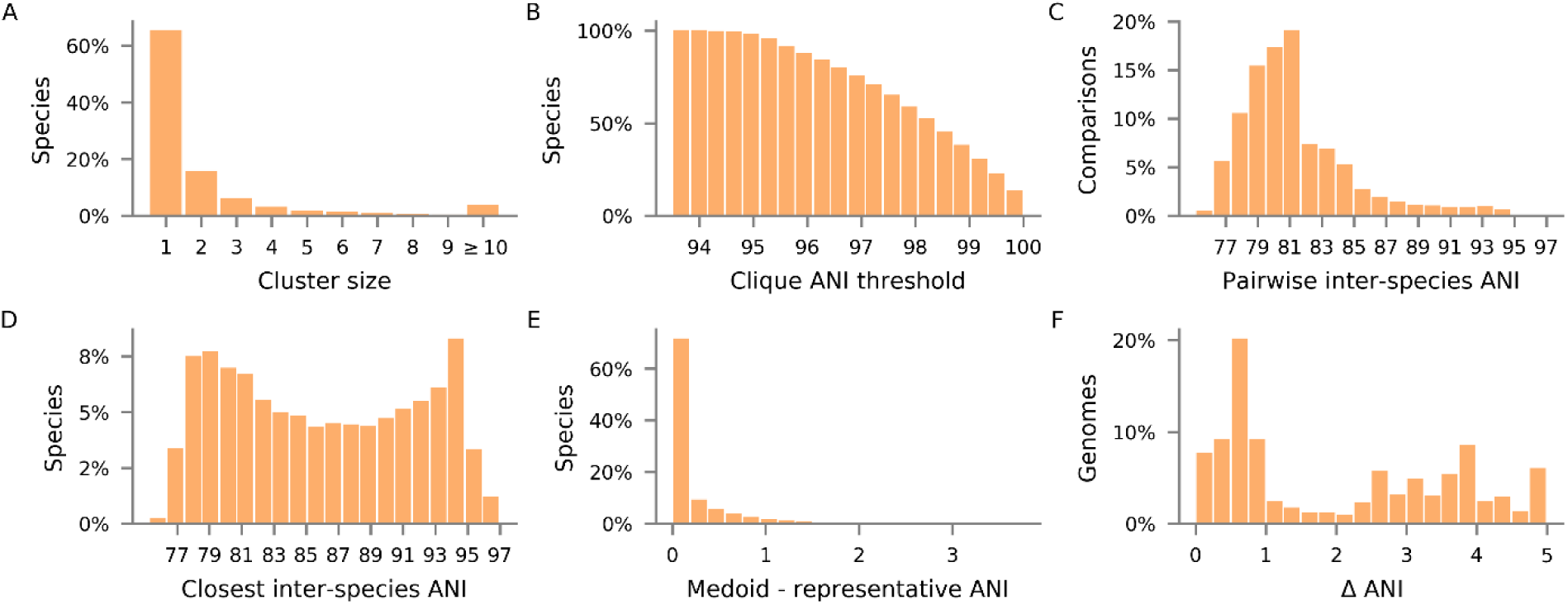
Key properties of GTDB species clusters circumscribed by ANI to a representative genome. (A) Number of genomes within each of the 24,706 species clusters. (B) Percentage of species forming a clique at varying ANI thresholds for the 4,660 species clusters with ≥3 genomes. (C) ANI values for 881,840 pairs of species representatives within the same genus. ANI values could not be calculated for 45,224 genome pairs due to insufficient homologous regions between the genomes. (D) ANI values between 19,466 species representatives and the closest representative within the same genus. For 432 species the closest representative had insufficient homology for a reliable ANI value to be calculated. (E) Difference in the mean ANI to the medoid and mean ANI to the selected representative for 4,660 species with ≥3 genomes. (F) Difference in ANI between the closest and second closest representative genome for the 26,129 genomes within the ANI circumscription radius of multiple species.

The tight intra-species ANI clustering combined with relatively small inter-species ANI values suggest that the genome used to represent a species cluster is not critical. This can be illustrated by considering the consequences of using the medoid genome within each of the 4,660 species clusters with ≥3 genomes as the representative, analogous to the approach taken by MiGA (Rodriguez-R et al., 2018). For 1,574 clusters, the medoid genome is the same as the representative genome proposed in this study. For the remaining 3,086 species clusters, we calculated the mean ANI from all genomes in the cluster to the medoid and proposed representative genomes. The difference between these mean ANI values was 0.35 ± 0.51% on average with the 90th percentile being 0.96% and the maximum difference being 3.76% (**Fig. 4E**; **Supp. Table 9**). Reassigning genomes within species clusters with ≥3 genomes with the medoid genomes acting as the species representative resulted in 121,445 (99.6%) of 121,939 genomes being assigned to the same species with 362 (0.3%) assigned to a different species and 132 (0.1%) failing the criteria for species assignment.

We examined the uniqueness of species assignments by considering the number of species containing genomes within the ANI circumscription radius of other species. 456 (5.3%) of the 8,579 species containing ≥2 genomes (i.e. at least one genome other than the representative genome) had genomes within the ANI radius of ≥2 species (0.90% ≥3 species; 0.23% ≥4 species). The 456 species are from both named (202) and *de novo* (254) species clusters, with over half (278) of these species having an ANI circumscription radius of 95%. While this indicates there can be ambiguity in species assignments, the average difference in ANI between the closest and second closest representative genome was relatively high at 2.0±1.6 (**Fig. 4F**) demonstrating that assigning genomes to the closest representative is robust. Notably, 26,129 (21.5%) of the 121,198 non-representative genomes are within the ANI circumscription radii of multiple species with 21,915 (83.9%) being from species in just four intensely classified clinically important genera: *Escherichia* with 13,131 genomes, *Salmonella* with 5,808 genomes, *Listeria* with 1,775 genomes, and *Neisseria* with 1,201 genomes (**Supp. Table 10**).

### Evaluating robustness of proposed species clusters

We further assessed the robustness of the proposed species clusters by forming new clusters that did not take into account type material or genome quality. The genome dataset was first simplified by randomly subsampling GTDB species clusters comprising >10 genomes to 10 genomes to reduce computational requirements and more evenly weight individual species. The resulting 49,902 genomes were organized into species clusters by randomly selecting genomes to act as species representatives until all genomes were assigned to a cluster as determined using the same clustering criteria as described for *de novo* species clusters (*see Methods*). Five independent trials were conducted in order to explore the impact of using randomly selected representatives to form species clusters (**Supp. Table 11**). In all trials, randomly seeded clusters were highly congruent with the proposed GTDB clusters with 98.3 ± 0.05% of the 24,706 clusters being identical and 99.4 ± 0.06% of the 49,902 genomes retaining the same species assignment. There were 129 proposed species clusters that were incongruent across all five random trials with the majority of these (85 species; 65.9%) having an ANI circumscription radius >95% as a result of being from highly sampled genera such as *Neisseria* (4 species), *Paenibacillus* (4 species), *Streptococcus* (4 species), and 16 *Streptomyces* (16 species; **Supp. Table 12**).

### Monophyly of proposed species clusters in concatenated protein trees

Here we assess the monophyly of 7,293 proposed species clusters belonging to 2,029 genera by inferring maximum-likelihood trees for each genus using a randomly selected outgroup from the sister lineage in the bacterial or archaeal GTDB reference tree. Only genera that could result in polyphyly were included in the analysis (i.e. comprised of >2 species with at least one species circumscribing ≥2 genomes). As before, species consisting of >10 genomes were randomly sampled to 10 genomes. Of the 7,293 species, 6,854 (94.0%) were recovered as monophyletic and 29,115 of 29,564 (98.5%) genomes had a species assignment congruent with the topology of the inferred trees (**Supp. Table 13**). Of the 2,592 genomes belonging to the 439 polyphyletic species, 2,143 (81.8%) had species assignments congruent with the topology of the tree. This indicates that polyphyly is the result of a small number of incongruent genomes as evidenced by 263 (59.9%) species having only one incorrectly placed genome in the genus tree (**Supp. Table 13**). For comparison, across the same set of trees there were 2,894 species defined by the NCBI taxonomy consisting of >2 genomes with 2,027 (70.0%) forming monophyletic groups. Of the 6,493 genomes assigned to one of the 867 polyphyletic NCBI species, 5,594 (86.2%) had species assignments congruent with the topology of the tree.

### Comparison to NCBI species assignments

Of the 143,566 genomes with an NCBI taxonomic assignment, 79.6% had a species assignment as a consequence of deeply sequenced species having recognized species names within the NCBI taxonomy (e.g. *Staphylococcus aureus* with 9,444 genome assemblies). Of these, 30.8% have a different species assignment under the proposed GTDB species clusters (**Fig. 5**). This is largely the result of reassignment of *Escherichia coli* genomes (26.7% of changes) to *E. flexneri* and *E. dysenteriae* along with changes to the generic name of a few deeply sampled species such as *Bacillus* (7.5%), *Campylobacter* (6.4%), and *Shigella* (5.1%; **Supp. Tables 14** and **15**). Arguably, the more relevant metric is only comparing species representatives to remove the distorting effect of highly sampled species. Using this criterion, less than half (10,323; 43%) of the 24,080 proposed species representatives with an NCBI taxonomic assignment were classified to the species level (**Fig. 5**). Of the NCBI-classified representatives, over a third (3,620; 35.1%) have a different species assignment in the GTDB (**Supp. Table 16**). The majority of these changes (2,473; 68.3%) were due to modifying the generic name of the species as a result of resolving polyphyletic genera or normalizing genera by relative evolutionary divergence (Parks et al., 2018). The 10 most commonly reassigned genus names account for >30% of the total differences between the two taxonomies (**Supp. Table 17**) and includes commonly recognized polyphyletic groups such as *Pseudomonas* (8.5%; Peix et al., 2009), *Bacillus* (5.5%, Bhandari et al., 2013), and *Clostridum* (3.3%; Beiko, 2015).

**Figure 5.**
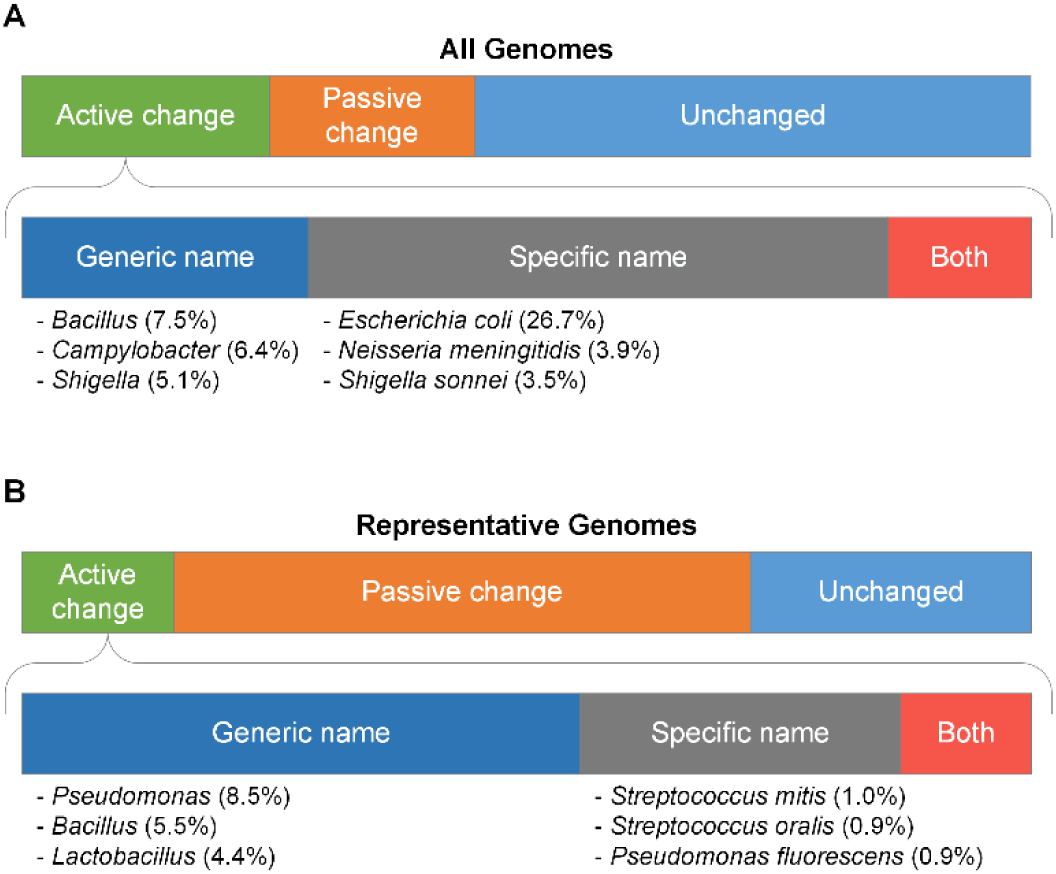
Comparison of proposed species assignments with the NCBI taxonomy. Results are shown for the 143,566 genomes (A) and 24,080 species representatives (B) with an NCBI taxonomic assignment. A genome was categorised as unchanged if its binomial species name was identical in the NCBI taxonomy, passively changed if the NCBI taxonomy did not have a species assignment, or actively changed if the proposed species name differs from the NCBI classification. In the lower bars, active changes were divided into those with a change in only the generic name of the species, only the specific name of the species, or due to changes in both the generic and specific names. The top three most commonly changed NCBI genera and species are listed.

### Genomic characteristics of bacterial and archaeal species

Organization of genomes into species clusters provides the opportunity to compare the distribution of common genomic properties across species representatives, which may be expected to differ substantially from distributions calculated over all genomes (**Fig. 6**; **Supp. Table 18**). For example, the genome size of bacterial and archaeal species representatives forms a skewed normal distribution in contrast to the binomial distribution which occurs when genome size is calculated over all genomes. This difference can be primarily attributed to over-representation of human pathogen species in the NCBI Assembly database (**Supp. Table 3**) and highlights the importance of calculating genome statistics over species rather than genomes.

**Figure 6.**
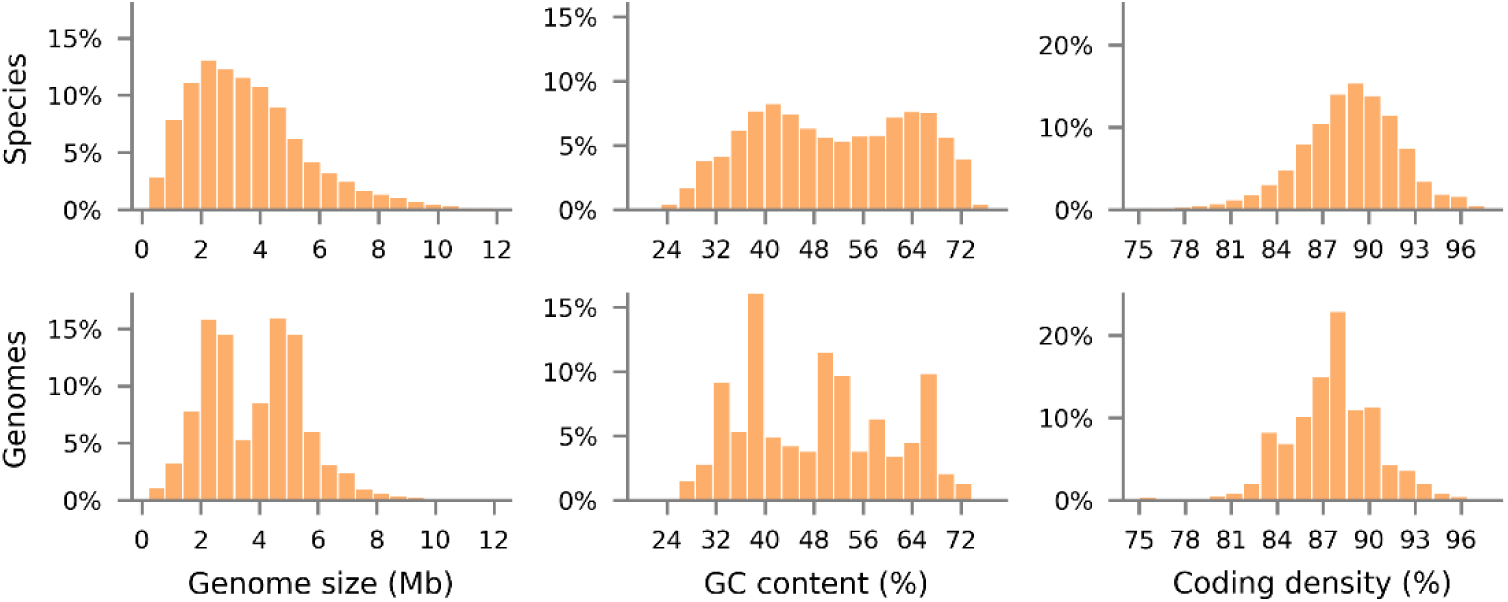
Distribution of common genomic parameters for the 24,706 species representative genomes and all 145,904 genomes in GTDB R04-RS89.

## Discussion

Type material is the cornerstone of modern prokaryote nomenclature and forms the basis for taxonomic opinion (Parker et al., 2019). Ideally, therefore, a genome-based taxonomy should be based on type material providing a reference point for both naming and classification. Here, we have taken this concept and combined it with ANI to produce a fully articulated species classification for publicly available bacterial and archaeal genomes. The proposed species clusters are circumscribed by ANI to a representative genome for each species using a genome assembled from the type strain of the species where possible. Identifying genomes assembled from type material is a challenge given the current practice of referencing type strains through co-identical identifiers (Federhen, 2015; Patterson et al., 2016; Salvà-Serra et al., 2019; Wittouck et al., 2019). This is highlighted by the 478 genomes for which only LPSN or BacDive, but not both, indicate a genome is assembled from the type strain of the species (**Fig. 1B**). Ideally, type material should be referenced by a single universal identifier which would be used as the primary identifier across all nomenclatural resources and genomic repositories.

Circumscribing species using ANI to representative genomes provides a quantitative and convenient operational species definition. However, this definition is not always congruent with species whose names are validly or effectively published as evident from 370 species becoming synonyms under the proposed species clusters (**Supp. Table 6)**. This is exemplified by the nine species in *Brucella* being reclassified into a single species, *Brucella melitensis*. While the genomic evidence supporting this reassignment has been recognized for over three decades (Verger et al., 1985), concerns have been raised in regards to the challenges this change would cause clinicians and regulatory agencies (Osterman and Moriyon, 2006; Gargani and Lopez-Merino 2006; Ficht 2010; Fenwick and Carroll 2019). We appreciate these concerns but have opted for the proposed quantitative species definition as we believe it is of the greatest use to the majority of the scientific community and best reflects current opinions regarding the circumscription of species. The term variant or subspecies will be incorporated into future GTDB releases to accommodate preserving historical nomenclature associations through reclassification of specific names as infrasubspecific epithets e.g., *Brucella vulpis* will be classified as *Brucella melitensis* var. *vulpis*, as previously proposed (Riojas et al., 2018).

Arguably the most contentious reclassifications resulting from the proposed species clusters are within the *Escherichia/Shigella* genus. GTDB classifies *Shigella* species as belonging to the genus *Escherichia* (Parks et al., 2018; **Supp. Table 15**), and *E. sonnei* and *E. boydii* are later heterotypic synonyms of *E. flexneri* under the proposed species clusters (**Supp. Table 6**). Furthermore, circumscription of species based on ANI to the type strains of *E. coli, E. flexneri*, and *E. dysenteriae* resulted in 7,212 and 1,183 genomes classified as *E. coli* within the NCBI taxonomy being reassigned to *E. flexneri* and *E. dysenteriae*, respectively (**Supp. Table 14**). This represents a reassignment of nearly 80% of the genomes classified as *E. coli* at NCBI. The scale of these reassignments results in traditional properties of these species no longer holding true, e.g., *E. dysenteriae* and *E. flexneri* being comprised of human pathogenic strains (Lan and Reeves, 2002). Consequently, it may be prudent to reassign *E. flexneri* and *E. dysenteriae* as synonyms of *E. coli* (96.4% and 96.2% ANI, respectively) to avoid confusion and to better reflect the high genomic similarity and evolutionary relationships of these species (Lan and Reeves, 2002). This change is being considered for the next release of the GTDB as an exception to defining synonyms at >97% ANI.

Nearly all proposed species clusters (98.2% of species) have an ANI circumscription radius of 95%. Consequently, we might expect intra-species ANI values to approach 90% reflecting the “10% diameter” around representative genomes. In practice, all species clusters form a clique at 93.5% ANI and the vast majority (97.6%) form a clique at 95% ANI. This tight intra-species clustering of genomes may reflect evolutionary forces shaping speciation since it is trivial to bioinformatically produce species clusters with a 10% diameter. Tight intra-species clustering has the practical benefit that the selection of species representatives is not critical as demonstrated by medoid and random representative testing (**Supp. Table 11**). Therefore, selection of type strains to represent species clusters is the most pragmatic choice since they are directly tied to nomenclature. The inter-species ANI between closest representatives within a genus were nearly evenly distributed between 78 and 95% ANI (**Fig. 4D**). This result is in contrast to reports of a genetic discontinuum between 83% to 95% ANI (Jain et al., 2018). These apparently contradictory results may reflect differences in how species were defined, but is primarily the result of changing the perspective from all pairwise comparisons in a large genomic dataset (Jain et al., 2018) to the inter-genus ANI values between closest species representatives (this study; Kim et al., 2014). This latter perspective suggests a fairly smooth continuum of inter-species generic diversity within genera which, despite the results presented here, may ultimately challenge efforts to unambiguously define species boundaries using ANI as additional species are discovered (Hanage, 2013). In this event, other means for defining species may ultimately be more appropriate, such as recombination boundaries (Bobay and Ochman, 2017; Arevalo et al., 2019).

Incorporation of the proposed species clusters into the GTDB requires that they be updated biannually with each GTDB release (Parks et al., 2018). An emphasis will be placed on maintaining the selected representative genome for each species cluster between GTDB releases so they can serve as stable nomenclatural type material (Whitman, 2016; Konstantinidis et al., 2017; Chuvochina et al., 2019). However, this must be balanced with the desire to use high-quality genomes assembled from the type strain of the species as representatives and the incorporation of changing taxonomic opinion. It is also likely that the genome assemblies of some selected representatives will ultimately be found to be in error, either in terms of the assembly itself or the associated metadata (i.e. incorrect species or strain assignment). We anticipate that the selection of a single representative genome acting as the nomenclatural type for each species will place these genomes under heavier community scrutiny which will help uncover such issues. Ultimately, these issues mean that some changes to species clusters will occur with each update.

The proposed quantitative species definition allows for the scalable and automated assignment of genomes to species clusters. We have incorporated these species clusters into the GTDB and GTDB-Tk (Chaumeil et al., 2019), an open source tool for the taxonomic classification of genome assemblies. These clusters encompass the nearly 150,000 public genomes within the NCBI Assembly database and will be updated with each GTDB release. This provides a complete genome-based taxonomy where all genomes have an assignment from domain to species and establishes representative genomes for circumscribing species. We anticipate that the availability of quantitatively defined and regularly updated species clusters will greatly improve the taxonomic resolution of microbial studies and consequently the communication of scientific results.

## Methods and Material

### Genome dataset

A dataset of 152,288 bacterial and 2,661 archaeal genomes was obtained from the NCBI Assembly database (Kitts et al., 2016) on 2018 July 16, and supplemented with 153 archaeal MAGs recovered from Sequence Read Archive metagenomes (Leinonen et al., 2011; Parks et al., 2018). These genomes comprise the bases of the Genome Taxonomy Database (GTDB) R04-RS89. Genomes were flagged for exclusion if they failed any of the following quality control criteria: i) estimated completeness <50%, ii) estimated contamination >10%, iii) completeness-5×contamination <50, iv) comprised of >1,000 contigs, v) N50 <5 kb, vi) contained >100,000 ambiguous bases, or vii) contained <40% of the 120 bacterial or 122 archaeal proteins used for phylogenetic inference. CheckM v1.0.13 (Parks et al., 2015) was used to estimate genome quality and determine assembly statistics. This filtering resulted in 8,185 genomes being flagged for removal with 87 being retained after manual inspection as they represent genomes of high nomenclatural or taxonomic significance (**Supp. Table 19**), *e.g.* the isolate genome *Ktedonobacter racemifer* representing the class Ktedonobacteria was retained despite having a contamination estimate of 11%.

### Genome metadata and standing in nomenclature

The NCBI taxonomy (Federhen, 2015) associated with each genome assembly was obtained from the NCBI FTP site on 2018 July 16. NCBI Assembly records for each genome were also downloaded on 2018 July 16 and used to establish if a genome was annotated as being assembled from type material at NCBI, considered a representative or reference genome at NCBI, and whether the assembly is a MAG, SAG, or isolate. LPSN (Parte 2018; consulted 16 April 2019), BacDive (Reimer, 2019; consulted 16 April 2019), and StrainInfo (Verslyppe et al., 2014; consulted 19 November 2018) were cross referenced with the strain identifiers at NCBI to establish which genomes were assembled from the type strain of a species or type strain of a subspecies. NCBI strain identifiers were obtained from the following fields: the infraspecific name and isolate fields in the assembly report, the strain and isolate fields in the genomic GenBank flat file, and the strain and isolate fields in the WGS master GenBank flat file. A genome was considered assembled from type material if any of its co-identical strain identifiers could be matched with a co-identical strain identifier at LPSN, BacDive, or StrainInfo. The NCBI metadata and nomenclatural status of each genome is provided in the GTDB R04-RS89 metadata file available from the GTDB website (https://gtdb.ecogenomic.org).

### Selecting representative genome for species clusters

A single representative genome was selected for each of the 9,162 validly or effectively published species names in the quality-controlled genome dataset (**Supp. Fig. 1)**. The majority of species (5,942 species; 64.9%) had a single genome in the dataset and this genome was selected as the representative of the species. Selection of representative genomes for the remaining 3,220 species were performed by prioritizing genomes in the following order: i) assembled from the type strain of the species (2,632 species), ii) annotated as being assembled from type material at NCBI (123 species), iii) designated as a reference or representative genome at NCBI (220 species), or iv) assembled from the type strain of a subspecies (8 species). Filtering by these metadata categories resulted in a further 1,714 species being reduced to a single genome that was selected as the representative of the species. Further selection criteria were used to establish a representative genome for the remaining 1,506 species which contained multiple genomes within a specific metadata category (e.g., multiple genomes from the type strain of the species). Of those, 196 species were resolved by selecting the single genome annotated as being assembled from type material at NCBI. A further 1,073 species were resolved by determining that all genomes formed a clique at ≥99% ANI and selecting the highest-quality genome. For this purpose, genome quality was defined with regards to multiple genome assembly statistics and metadata fields indicative of assembly quality (**Supp. Table 20**). From the remaining 237 species that had at least one genome pair with an ANI <99%, 18 were resolved by selecting the single genome annotated as a representative or reference genome at NCBI. The final 219 species were resolved manually through investigation of the literature, inspection of 16S rRNA homology results, and consideration of all pairwise ANI results within a species.

### Calculating average nucleotide identity and alignment fraction

The ANI and AF between genomes was calculated using FastANI v1.1 (Jain et al., 2018) with default parameters. FastANI requires at least 150 Kb of homologous genome sequence between two genomes to make a reliable ANI estimate. ANI and AF values are not symmetric and were defined as the maximum of the two possible values. Using the maximum AF accommodates MAGs, SAGs, and lower-quality isolate genomes that can be incomplete or contaminated. Use of the maximum ANI value is for convenience and is not a critical decision as ANI values are nearly symmetrical in practice (Jain et al., 2018).

### Circumscribing named species clusters

Clusters for validly or effectively named species were established by i) determining the species-specific ANI circumscription radius for each representative genome, and ii) assigning non-representative genomes to species clusters based on these ANI thresholds. The ANI circumscription radius for a representative genome was determined by calculating the ANI to all other representatives and setting the ANI circumscription radius as follows: i) 95% if the closest representative had an ANI <95% ANI or ii) to the ANI value of the closest representative if this was between 95% and 97%. Any representative with an ANI >97% was considered a synonym and disregarded for the purposes of establishing the ANI circumscription radius. To reduce computational requirements, Mash v2.1.1 (Ondov et al., 2016) was used to produce a subset of representative genome pairs with a Mash distance of ≤0.1 for processing by FastANI. The ANI and AF between non-representative and representative genomes were calculated using FastANI, again using Mash with a distance threshold of ≤0.1 as a prefilter. A non-representative genome was assigned to the closest representative genome where the AF was >65% only if it was within the ANI circumscription radius of the representative.

### Establishing de novo species clusters

Genomes not assigned to a named species cluster were formed into *de novo* species clusters using a greedy clustering approach which favored the selection of high-quality representative genomes. Genome quality was defined in the same manner used to resolve the selection of representative genomes for validly or effectively published species with multiple genomes assembles (**Supp. Table 20**). Greedy clustering consists of four steps: i) sort genomes without a species assignment by genome quality, ii) select the highest-quality genome to form a new species cluster, iii) determine the species-specific ANI circumscription radius for the new species cluster, and iv) assign genomes without a species assignment to this species cluster using the same criteria as for named species clusters. These steps are repeated until all genomes have been assigned to a species. Determining the species-specific ANI circumscription radius (step 3) requires calculating the ANI to all existing species representatives and setting the ANI radius to 95% if the closest representative had an ANI <95% ANI or to the ANI value of the closest representative. The ANI to the closest representative will always be ≤97% by definition and thus synonyms need not be considered during the formation of *de nov*o species clusters. This procedure ensures that all genomes can be assigned to at least one species representative, but does not guarantee that all genomes are assigned to the closest representative. Consequently, all non-representative genomes were reassigned to the closest representative where the AF is >65% and where the genome is within the ANI circumscription radius of the representative.

### Naming de novo species clusters

Species clusters consisting of genomes without a validly or effectively published name were assigned a placeholder name. The generic name follows the genus name within the GTDB which is established as described previously (Parks et al., 2018). The specific name was derived from the NCBI accession of the representative genome of the species cluster. Specifically, the numerical portion of the accession is prefixed with ‘sp’. For example, GCF_000192635.1 is the representative genome of a species cluster within the genus Agrobacterium so was assigned the species name Agrobacterium sp000192635.

Genera and species names with an alphabetic suffix indicate genera and species that are polyphyletic or needed to be subdivided based on taxonomic rank normalisation according to the current GTDB reference tree. The lineage containing the type strain retains the unsuffixed name and all other lineages are given alphabetic suffixes, indicating that they are placeholder names that need to be replaced in due course. For example, the proposed species clusters define both *Lactobacillus gasseri* and Lactobacillus gasseri_A, the latter being a *de novo* GTDB species cluster which does not contain the *L. gasseri* type strain and circumscribes one or more genomes classified as *L. gasseri* in the NCBI taxonomy.

### Properties of species clusters

Species clusters containing >100 genomes were randomly sampled to 100 genomes. Inter-species properties were calculated by limiting the calculation to species within the same genus and discarding comparisons with an AF <25%. The medoid genome of a species clusters is defined as the genome minimizing the mean ANI to all genomes in the cluster and was determined using a brute-force implementation of this definition. For species clusters with 2 genomes, the medoid was taken as the selected representative genome as this genome is of higher quality.

### Assessing robustness of species clusters using randomly selected representatives

Species clusters consisting of >10 genomes were randomly sampled to 9 genomes along with the proposed representative genome. Random species representatives were selected in a manner similar to that described for the *de novo* species clusters: i) randomly select a genome without a species assignment, ii) set the species-specific ANI circumscription radius to the radius of its proposed species cluster, and iii) assign genomes without a species assignment to this species cluster using the same criteria as for named species clusters. These steps are repeated until all genomes have been assigned to a species. All non-representative genomes were then reassigned to the closest representative where the AF is >65% and where the genome is within the ANI circumscription radius of the representative. This procedure was independently repeated 5 times. The proposed and random species clusters were compared to determine i) the number of clusters in perfect congruence (i.e., there exists a proposed and random species cluster comprised of the exact same set of genomes) and ii) the number of genomes that would be assigned the same species name (i.e. the number of genomes in common between each proposed species cluster and the random species cluster containing the representative genome of the proposed species cluster).

### Assessing monophyly of species clusters

The monophyly of the proposed species clusters was assessed by inferring trees for each genus where polyphyly can arise (i.e. genera comprised of ≥2 species with at least one species containing ≥2 genomes). Species clusters consisting of >10 genomes were randomly sampled to 10 genomes. Trees were rooted by randomly selecting a genome from the sister lineage to the genus as determined from the topology of the GTDB R04-RS89 reference trees. The GTDB-Tk v0.3.2 (Chaumeil et al., 2019) ‘de novo’ workflow with default parameters was used to infer the trees. Briefly, genes were called using Prodigal v2.6.3 (Hyatt et al., 2010) and 120 bacterial or 122 archaeal marker genes identified and aligned using HMMER v3.1b (Eddy, 2011). The resulting multiple sequence alignment was trimmed to approximately 5,000 columns using the bacterial or archaeal GTDB R04-RS89 mask. Trees were inferred with FastTree v2.1.10 (Price et al., 2009) with the WAG+GAMMA models and rooted on the selected outgroup. PhyloRank v0.0.37 (https://github.com/dparks1134/phyloRank) was used to decorate the tree with the proposed species assignments and determine the F-measure for each species (McDonald et al., 2012). A genome was considered to be congruent with the topology of the tree if it was contained in the lineage with the highest F-measure for its corresponding species assignment.

### Code availability

The methodology used to establish the GTDB species clusters is implemented in the GTDB Species Cluster Toolkit (https://github.com/Ecogenomics/gtdb-species-clusters), a Python program available under the GNU General Public License v3.0. The code used to establish the species clusters in GTDB R04-RS89 is retained under version GTDB-R89.

### Data availability

The metadata associated with each genome and used to establish the GTDB species clusters is available on the GTDB website (http://gtdb.ecogenomic.org) in the files ar122_metadata_r89.tsv and bac120_metadata_r89.tsv. The genomic files for genomes are available from the NCBI Assembly database, including the 153 archaeal MAGs which are under BioProject <in_submission>.

## Acknowledgements

We thank the members of the Microbial Taxonomy Workshop for helpful discussions relating to establishing species clusters. This project is primarily supported by an Australian Research Council Laureate Fellowship (FL150100038) awarded to PH.

## Author contributions

D.P. and P.H. wrote the paper with constructive suggestions from all other authors. D.P. designed the initial study. M.C., C.R., and P.H. provided nomenclatural advice and manual curation of species representatives where necessary. D.P., P-A.C., and A.M. performed the bioinformatic analyses.

