## Supplemental figures and tables for "Selection of representative genomes for 24,706 bacterial and archaeal species clusters provide a complete genome-based taxonomy"

### Supplementary Figures

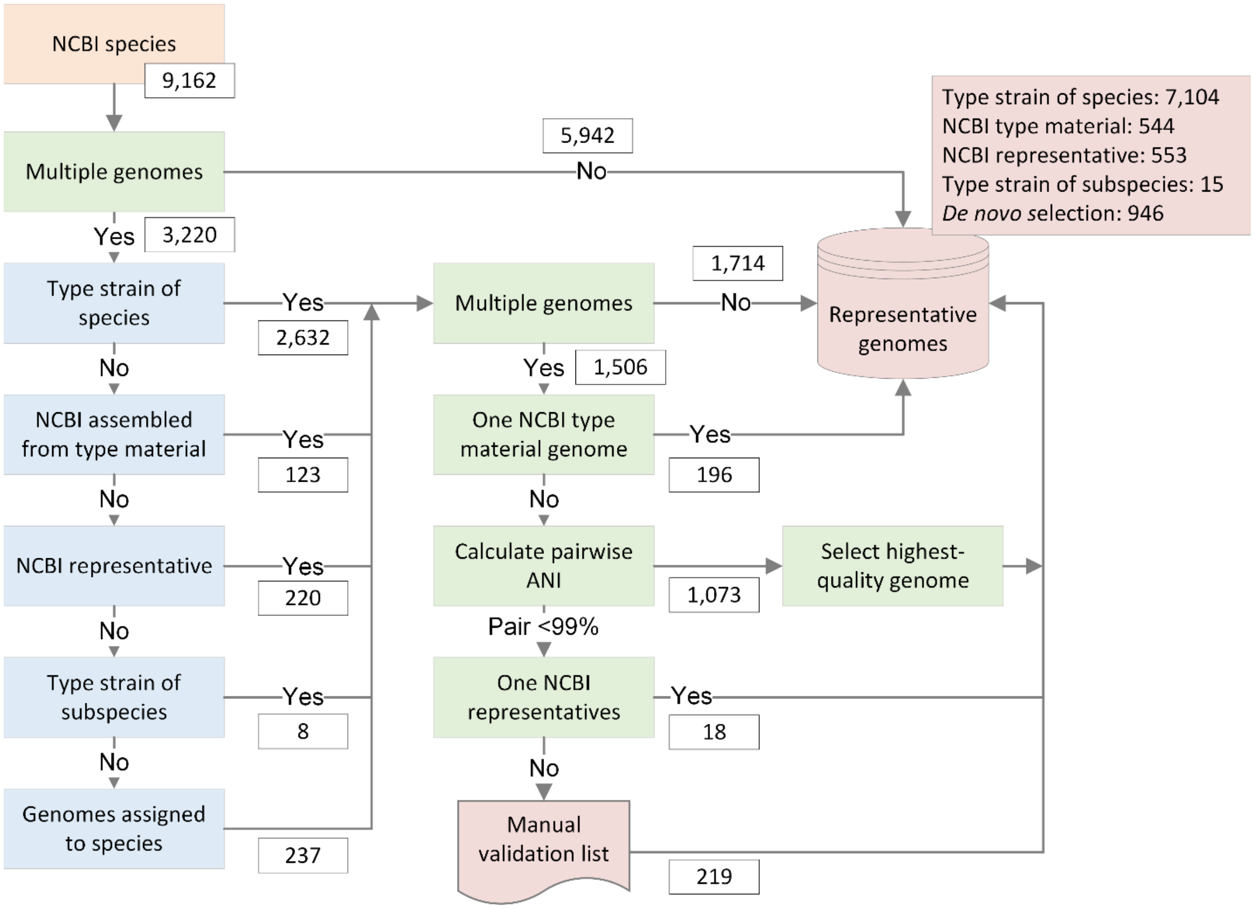


**Supp. Fig. 1**. Workflow for selecting a single genome representative for each validly or effectively published species. Numbers of genomes following different paths through the workflow are indicated in rectangles.

### Supplementary Tables

**Supp. Table 1**. Number of genomes and species identified as being assembled from type material.

| | Genomes | | | Species | | |
|---|---|---|---|---|---|---|
| | *Bacteria* | *Archaea* | *Total* | *Bacteria* | *Archaea* | *Total* |
| Type strain of species | 8,313 | 352 | 8,665 | 6,794 | 310 | 7,104 |
| Type strain of subspecies | 146 | 0 | 146 | 83 | 0 | 83 |

**Supp. Table 2**. Percentage of the 9,162 species representative genomes assigned to different metadata categories.

| Category | Bacteria and Archaea | Bacteria | Archaea |
|---|---|---|---|
| No. validly or effectively published species | 9,162 | 8,764 | 398 |
| Type strain of species | 7,104 (77.53%) | 6,794 (77.52%) | 310 (77.89%) |
| NCBI type material | 544 (5.94%) | 531 (6.06%) | 13 (3.27%) |
| NCBI representative | 553 (6.04%) | 535 (6.10%) | 18 (4.52%) |
| Other | 961 (10.49%) | 904 (10.31%) | 57 (14.32%) |
| Single genome in cluster | 5,942 (64.85%) | 5,633 (64.27%) | 309 (77.64%) |
| Multiple genomes in cluster | 3,220 (35.15%) | 3,131(35.73%) | 89 (22.36%) |
| Single nomenclatural source | 393 (4.29%) | 385 (4.39%) | 8 (2.01%) |
| Multiple nomenclatural sources | 6,711 (73.25%) | 6,409 (73.13%) | 302 (75.88%) |
| No nomenclatural sources | 2,058 (22.46%) | 1,970 (22.48%) | 88 (22.11%) |

**Supp. Table 3.** Species with the largest number of genomes in GTDB R04-RS89.

| Species | No. genomes |
|---|---|
| *Staphylococcus aureus* | 9,444 |
| *Escherichia flexneri* | 9,084 |
| *Salmonella enterica* | 8,698 |
| *Streptococcus pneumoniae* | 8,201 |
| *Mycobacterium tuberculosis* | 5,596 |
| *Klebsiella pneumoniae* | 4,086 |
| *Acinetobacter baumannii* | 2,796 |
| *Pseudomonas aeruginosa* | 2,744 |
| *Escherichia coli* | 2,432 |
| *Mycobacteroides abscessus* | 1,589 |

**Supp. Table 4.** Genomes satisfying the assignment criteria of multiple species with validly or effectively published names (see Excel file).

**Supp. Table 5**. Number of validly or effectively published species names that become synonyms for varying ANI thresholds.

| ANI synonym criterion | No. validly published synonyms | No. effectively published synonyms |
|---|---|---|
| 99.5% | 39 | 38 |
| 99% | 60 | 85 |
| 98% | 126 | 155 |
| 97% | 176 | 194 |
| 96% | 224 | 235 |

**Supp. Table 6.** Species reclassified as synonyms because they have an ANI >97% to a species with naming priority (see Excel file).

**Supp. Table 7**. Percentage of the 24,706 GTDB representative genomes assigned to different metadata categories.

| Category | Bacteria and Archaea | Bacteria | Archaea |
|---|---|---|---|
| Total species clusters | 24,706 | 23,458 | 1,248 |
| Named species clusters | 8,792 (35.59%) | 8,412 (35.86%) | 380 (30.45%) |
| *De novo* species clusters | 15,914 (64.41%) | 15,046 (64.14%) | 868 (69.55%) |
| MAG | 9,861 (39.91%) | 9,086 (38.73%) | 775 (62.10%) |
| SAG | 439 (1.78%) | 416 (1.77%) | 23 (1.84%) |
| Isolate | 14,406 (58.31%) | 13,956 (59.49%) | 450 (36.06%) |
| Single genome in cluster | 16,127 (65.28%) | 15,264 (65.07%) | 863 (69.15%) |
| Multiple genomes in cluster | 8,579 (34.72%) | 8,194 (34.93%) | 385 (30.85%) |

**Supp. Table 8**. Statistics for species clusters.

| **Statistic (species clusters with > 2 genomes)** | **Named species** | ***De novo* species** | **All species** |
|---|---|---|---|
| No. species clusters | 2,148 | 2,512 | 4,660 |
| Mean intra-species pairwise ANI | 98.4 ± 1.03 | 98.8 ± 1.13 | 98.6 ± 1.10 |
| Mean intra-species ANI to representative genome | 98.4 ± 1.08 | 98.8 ± 1.21 | 98.6 ± 1.16 |
| Mean intra-species ANI to medoid – ANI to representative | 0.35 ± 0.48 | 0.35 ± 0.54 | 0.35 ± 0.51 |
| Clique at 95% ANI | 2,084 (97.0%) | 2,450 (97.5%) | 4,534 (97.3%) |
| Clique at 94% ANI | 2,143 (99.8%) | 2,511 (100.0%) | 4,654 (99.9%) |
| Minimum intra-species ANI value | 93.5% | 93.9% | 93.5% |
| ANI circumscription radius = 95% | 1,987 (92.5%) | 2,493 (99.2%) | 4,480 (96.1%) |

| **Statistic (species clusters with ≥ 2 genomes)** | **Named species** | ***De novo* species** | **All species** |
|---|---|---|---|
| No. species clusters | 3,618 | 4,961 | 8,579 |
| Mean intra-species pairwise ANI | 98.6 ± 1.20 | 98.8 ± 1.22 | 98.7 ± 1.21 |
| Mean intra-species ANI to representative genome | 98.6 ± 1.22 | 98.8 ± 1.26 | 98.7 ± 1.25 |
| Clique at 95% ANI | 3,554 (98.2%) | 4,899 (98.8%) | 8,453 (98.5%) |
| Clique at 94% ANI | 3,613 (99.9%) | 4,960 (100.0%) | 8.573 (99.9%) |
| Minimum intra-species ANI value | 93.5% | 93.9% | 93.5% |
| ANI circumscription radius = 95% | 3,397 (93.9%) | 4,929 (99.5%) | 8.326 (97.1%) |

| **Statistics (all species clusters)** | **Named species** | ***De novo* species** | **All species** |
|---|---|---|---|
| No. species clusters | 8,792 | 15,914 | 24,706 |
| ANI circumscription radius = 95% | 8,407 (95.6%) | 15,853 (99.6%) | 24,260 (98.2%) |

**Supp. Table 9.** Species where the difference in the mean ANI from the medoid and mean ANI from the GTDB-selected representative is >2 (see Excel file).

**Supp. Table 10.** Genomes satisfying the assignment criteria for multiple GTDB species clusters (see Excel file).

**Supp. Table 11**. Comparison of species clusters formed from randomly selected representative genomes to the proposed species clusters.

| **Species clusters** | **No. species clusters** | **Identical clusters** | **Identical clusters with ≥2 genome** | **Genomes with same species assignment** |
|---|---|---|---|---|
| *Proposed* | 24,706 | - | 8,579 | - |
| Random 1 | 24,696 | 24,310 (98.4%) | 8,273 (96.4%) | 49,629 (99.5%) |
| Random 2 | 24,690 | 24,271 (98.2%) | 8,246 (96.1%) | 49,573 (99.3%) |
| Random 3 | 24,714 | 24,295 (98.3%) | 8,265 (96.3%) | 49,583 (99.4%) |
| Random 4 | 24,675 | 24,299 (98.4%) | 8,262 (96.3%) | 49,644 (99.5%) |
| Random 5 | 24,674 | 24,297 (98.3%) | 8,268 (96.4%) | 49,641 (99.5%) |
| *Average ± Std* | 24,690 ± 16.5 | 98.3 ± 0.05 | 96.3 ± 0.11 | 99.4 ± 0.06 |

**Supp. Table 12.** Species clusters that were incongruent with the proposed species clusters across all five trials evaluating the impact of forming species clusters from randomly selected representative genomes (see Excel file).

**Supp. Table 13.** Evaluation of the monophyly of the proposed species clusters (see Excel file).

**Supp. Table 14.** Most commonly reassigned species between the NCBI and GTDB taxonomies across all genomes with a classification in the NCBI taxonomy. Numbers following GTDB species names indicate the number of genomes affected.

| NCBI species | No. reassigned genomes | Reassigned genomes (%) | GTDB assignments |
|---|---|---|---|
| *Escherichia coli* | 9,416 | 26.7 | Escherichia flexneri:7212; Escherichia dysenteriae:1183; Escherichia coli_D:974; Escherichia coli_C:27; Escherichia sp000208585:7; Escherichia marmotae:4; Klebsiella pneumoniae:2; Citrobacter freundii:2; Escherichia albertii:1; Klebsiella_B aerogenes:1; Klebsiella quasipneumoniae:1; Klebsiella variicola:1; Enterobacter himalayensis:1 |
| *Neisseria meningitidis* | 1,386 | 3.9 | Neisseria meningitidis_B:1374; Neisseria subflava_B:4; Neisseria sp000186165:3; Neisseria meningitidis_A:2; Neisseria flavescens:2; Neisseria sp000227275:1 |
| *Shigella sonnei* | 1,248 | 3.5 | Escherichia flexneri:1241; Escherichia coli_D:6; Serratia marcescens_I:1 |
| *Campylobacter jejuni* | 1,226 | 3.5 | Campylobacter_D jejuni:1211; Campylobacter_D coli:9; Campylobacter_D lari_C:3; Campylobacter_D jejuni_C:1; Campylobacter_D jejuni_A:1; Campylobacter_D jejuni_B:1 |
| *Listeria monocytogenes* | 1,157 | 3.3 | Listeria monocytogenes_B:1084; Listeria monocytogenes_C:67; Listeria innocua:4; Enterococcus faecalis:1; Enterococcus_B thailandicus:1 |
| *Enterococcus faecium* | 1,016 | 2.9 | Enterococcus_B faecium:913; Enterococcus_B faecium_B:97; Enterococcus faecalis:2; Enterococcus_D sp002850555:2; Enterococcus_B hirae:1; Enterococcus_B durans:1 |
| *Bacillus cereus* | 990 | 2.8 | Bacillus_A thuringiensis_J:379; Bacillus_A cereus_AD:151; Bacillus_A cereus:98; Bacillus_A cereus_P:68; Bacillus_A anthracis:45; Bacillus_A wiedmannii:40; Bacillus_A cereus_T:29; Bacillus_A thuringiensis:24; Bacillus_A cereus_AX:23; Bacillus_A thuringiensis_N:19; Bacillus_A toyonensis:19; Bacillus_A mycoides:16; Bacillus_A thuringiensis_S:13; Bacillus_A cereus_U:9; Bacillus_A cereus_S:7; Bacillus_A cereus_K:7; Bacillus_A cereus_AT:5; Bacillus_A cereus_Q:5; Bacillus_A cereus_AU:5; Bacillus_A thuringiensis_K:4; Bacillus_A thuringiensis_M:4; Bacillus_A sp002584985:3; Bacillus_A sp001884105:3; Bacillus_A pseudomycoides:3; Bacillus_A cereus_AK:2; Bacillus_A cereus_AG:2; Bacillus_A cereus_AQ:2; Bacillus_A cereus_AW:1; Bacillus_A cereus_AV:1; Bacillus_A cereus_O:1; Bacillus_A cereus_AY:1; Bacillus_A mycoides_B:1 |
| *Campylobacter coli* | 813 | 2.3 | Campylobacter_D coli:700; Campylobacter_D coli_A:64; Campylobacter_D coli_B:46; Campylobacter_D jejuni:3 |
| *Burkholderia pseudomallei* | 738 | 2.1 | Burkholderia mallei:738 |
| *Enterobacter cloacae* | 677 | 1.9 | Enterobacter himalayensis:445; Enterobacter nimipressuralis:70; Enterobacter kobei:41; Enterobacter cloacae_M:24; Enterobacter bugandensis:21; Enterobacter ludwigii:19; Enterobacter cloacae_H:12; Enterobacter cloacae_B:9; Enterobacter cloacae_J:6; Enterobacter sesami:5; Enterobacter cloacae_L:5; Enterobacter cloacae_I:5; Enterobacter sp000568095:3; Klebsiella_B aerogenes:2; Klebsiella pneumoniae:2; Enterobacter sp000493015:2; Kosakonia cowanii:1; Enterobacter mori:1; Leclercia sp000755535:1; Enterobacter asburiae:1; Enterobacter cloacae_K:1; Citrobacter portucalensis:1 |

**Supp. Table 15.** Most commonly reassigned genera between the NCBI and GTDB taxonomies across all genomes with a classification in the NCBI taxonomy. Numbers following GTDB genus names indicate the number of genomes affected.

| NCBI genus | No. reassigned genomes | Reassigned genomes (%) | GTDB assignments |
|---|---|---|---|
| *Bacillus* | 2,636 | 7.5 | Bacillus_A:2244; Bacillus_C:108; Bacillus_E:29; Bacillus_H:27; Bacillus_X:21; Bacillus_W:17; Bacillus_F:16; Bacillus_AA:15; Bacillus_I:10; Bacillus_J:10; Bacillus_L:9; Bacillus_AC:8; Bacillus_U:8; Bacillus_N:8; Bacillus_Y:6; Bacillus_AW:5; Bacillus_Q:5; Bacillus_AR:4; Bacillus_G:4; Bacillus_K:4; Bacillus_AZ:3; Bacillus_Z:3; Virgibacillus_A:3; Bacillus_AY:3; Bacillus_O:3; Alkalicoccus:3; Bacillus_AD:3; Bacillus_S:3; Bacillus_T:3; Bacillus_B:3; Bacillus_AI:2; Bacillus_AU:2; Bacillus_AJ:2; Bacillus_P:2; Bacillus_AX:2; Bacillus_M:2; Bacillus_AS:2; Bacillus_BJ:2; Bacillus_AH:2; Salisediminibacterium:2; Bacillus_BH:2; Bacillus_BB:1; Bacillus_BE:1; Bacillus_BD:1; Bacillus_AL:1; Bacillus_BC:1; Bacillus_R:1; Bacillus_AG:1; Bacillus_AP:1; Bacillus_BF:1; Bacillus_BL:1; Bacillus_AO:1; Virgibacillus:1; Bacillus_AE:1; Bacillus_AB:1; Bacillus_V:1; Bacillus_AV:1; Enterobacter:1; Lysinibacillus_F:1; Bacillus_BK:1; Bacillus_AN:1; Bacillus_AM:1; Bacillus_BA:1; Paenibacillus_S:1; Bacillus_AT:1; Bacillus_AF:1; Bacillus_AK:1 |
| *Campylobacter* | 2,262 | 6.4 | Campylobacter_D:2066; Campylobacter_A:176; Campylobacter_B:19; Corynebacterium:1 |
| *Shigella* | 1,782 | 5.1 | Escherichia:1778; Serratia:4 |
| *Pseudomonas* | 1,475 | 4.2 | Pseudomonas_E:1137; Pseudomonas_A:252; Pseudomonas_B:32; Pseudomonas_D:17; Pseudomonas_F:9; Pseudomonas_H:4; Pseudomonas_K:4; Pseudomonas_M:3; Stenotrophomonas:3; Pseudomonas_O:2; Burkholderia:2; Pseudomonas_G:2; Pseudomonas_L:2; Paraburkholderia:1; UBA6156:1; Pseudomonas_C:1; Pseudomonas_N:1; Xanthomonas:1; Corynebacterium:1 |
| *Lactobacillus* | 1,297 | 3.7 | Lactobacillus_F:302; Lactobacillus_C:274; Lactobacillus_G:209; Lactobacillus_H:202; Lactobacillus_B:163; Lactobacillus_D:58; Lactobacillus_K:33; Lactobacillus_O:19; Lactobacillus_E:18; Lactobacillus_J:6; Lactobacillus_I:4; Lactobacillus_A:2; Lactobacillus_N:2; Lactobacillus_L:2; Lactobacillus_M:2; Lachnospira:1 |
| *Enterococcus* | 1,198 | 3.4 | Enterococcus_B:1086; Enterococcus_D:41; Enterococcus_A:25; Enterococcus_E:25; Enterococcus_C:9; Enterococcus_G:4; Enterococcus_F:3; Enterococcus_H:2; Pseudomonas_E:1; Enterococcus_I:1; Enterococcus_J:1 |
| *Clostridium* | 584 | 1.7 | Clostridium_F:187; Clostridium_P:104; Clostridium_H:54; Clostridium_M:35; Clostridium_B:25; Clostridium_G:16; Clostridium_S:15; Clostridium_J:11; Clostridium_I:11; Hungatella:10; Ruminiclostridium_A:9; Clostridium_AM:8; Absiella:8; Erysipelatoclostridium:7; Clostridium_Q:7; Ruminiclostridium_B:5; Clostridium_L:4; Ruminiclostridium_F:4; Clostridium_E:4; Clostridium_X:4; Clostridium_W:4; Dorea:3; Clostridium_R:3; Clostridium_A:3; Clostridium_AO:3; Clostridium_AA:2; Clostridium_AI:2; Clostridium_Y:2; Clostridium_V:2; Acetivibrio:2; Clostridium_AH:2; Clostridium_K:2; Clostridium_T:2; Clostridium_Z:2; Ruminiclostridium:1; Clostridium_AB:1; Faecalicatena:1; Clostridium_AD:1; Clostridium_AE:1; Ruminiclostridium_D:1; Romboutsia:1; Clostridium_N:1; Clostridium_AC:1; Intestinibacter:1; Massilioclostridium:1; Corynebacterium:1; Clostridium_AG:1; Clostridium_D:1; Clostridium_AN:1; Clostridium_C:1; Lawsonibacter:1; Clostridium_AK:1; Clostridium_AF:1; Ruminiclostridium_C:1; Tyzzerella:1; Clostridium_U:1 |
| *Klebsiella* | 338 | 1.0 | Klebsiella_A:178; Klebsiella_B:149; Enterobacter:4; Raoultella:3; Atlantibacter:1; Escherichia:1; Serratia:1; Cedecea:1 |
| *Mycoplasma* | 305 | 0.9 | Mycoplasmoides:121; Mycoplasmopsis:55; Metamycoplasma:41; Mesomycoplasma:36; Mycoplasmopsis_A:28; Eperythrozoon_A:8; Mycoplasmopsis_B:5; Eperythrozoon_B:4; Malacoplasma:3; Malacoplasma_A:2; Mycoplasma_J:1; Mycoplasma_H:1 |
| *Helicobacter* | 181 | 0.5 | Helicobacter_E:70; Helicobacter_C:44; Helicobacter_D:33; Helicobacter_A:22; Helicobacter_G:4; Helicobacter_B:4; Helicobacter_H:2; Helicobacter_F:2 |

**Supp. Table 16.** GTDB and NCBI species assignments for the 24,080 proposed representative genomes with a species classification in the NCBI taxonomy (see Excel file).

**Supp. Table 17.** Most commonly reassigned genera between the NCBI and GTDB taxonomies across the GTDB representative genomes. Numbers following GTDB genus names indicate the number of genomes affected.

| NCBI genus | No. reassigned representatives | Reassigned representatives (%) | GTDB genus assignments |
|---|---|---|---|
| *Pseudomonas* | 308 | 8.5 | Pseudomonas_E:235; Pseudomonas_A:33; Pseudomonas_D:13; Pseudomonas_F:6; Pseudomonas_B:6; Pseudomonas_K:4; Pseudomonas_H:1; Pseudomonas_O:1; Paraburkholderia:1; Pseudomonas_G:1; UBA6156:1; Stenotrophomonas:1; Pseudomonas_M:1; Pseudomonas_C:1; Pseudomonas_L:1; Pseudomonas_N:1; Burkholderia:1 |
| *Bacillus* | 199 | 5.5 | Bacillus_A:37; Bacillus_AA:13; Bacillus_W:13; Bacillus_X:11; Bacillus_F:8; Bacillus_H:7; Bacillus_L:5; Bacillus_Q:5; Bacillus_Y:5; Bacillus_E:4; Bacillus_AW:4; Bacillus_AR:4; Bacillus_G:4; Bacillus_C:4; Bacillus_AZ:3; Bacillus_AY:3; Alkalicoccus:3; Bacillus_AD:3; Bacillus_K:3; Bacillus_AC:2; Bacillus_AI:2; Bacillus_Z:2; Bacillus_AJ:2; Bacillus_J:2; Bacillus_AX:2; Bacillus_S:2; Bacillus_AS:2; Salisediminibacterium:2; Bacillus_U:2; Bacillus_I:2; Bacillus_N:2; Bacillus_B:2; Bacillus_BB:1; Bacillus_AU:1; Bacillus_BE:1; Virgibacillus_A:1; Bacillus_BD:1; Bacillus_P:1; Bacillus_AL:1; Bacillus_BC:1; Bacillus_R:1; Bacillus_AG:1; Bacillus_M:1; Bacillus_AP:1; Bacillus_BF:1; Bacillus_BL:1; Bacillus_O:1; Bacillus_T:1; Bacillus_AO:1; Virgibacillus:1; Bacillus_AE:1; Bacillus_AB:1; Bacillus_V:1; Bacillus_AV:1; Lysinibacillus_F:1; Bacillus_BK:1; Bacillus_BJ:1; Bacillus_AN:1; Bacillus_AH:1; Bacillus_BH:1; Bacillus_AM:1; Bacillus_BA:1; Paenibacillus_S:1; Bacillus_AT:1; Bacillus_AF:1; Bacillus_AK:1 |
| *Lactobacillus* | 160 | 4.4 | Lactobacillus_G:50; Lactobacillus_H:23; Lactobacillus_K:19; Lactobacillus_B:16; Lactobacillus_O:13; Lactobacillus_C:12; Lactobacillus_F:7; Lactobacillus_D:5; Lactobacillus_E:4; Lactobacillus_J:3; Lactobacillus_I:2; Lactobacillus_L:2; Lactobacillus_N:1; Lachnospira:1; Lactobacillus_M:1; Lactobacillus_A:1 |
| *Clostridium* | 118 | 3.3 | Ruminiclostridium_A:8; Hungatella:8; Clostridium_H:7; Clostridium_B:6; Clostridium_M:6; Clostridium_F:5; Clostridium_J:5; Clostridium_I:5; Clostridium_AM:4; Clostridium_E:4; Clostridium_S:3; Erysipelatoclostridium:3; Clostridium_P:3; Clostridium_AO:3; Clostridium_L:2; Clostridium_G:2; Dorea:2; Clostridium_AA:2; Clostridium_AI:2; Clostridium_R:2; Clostridium_Y:2; Clostridium_W:2; Clostridium_Q:2; Clostridium_Z:2; Ruminiclostridium:1; Clostridium_AB:1; Faecalicatena:1; Clostridium_AD:1; Clostridium_AE:1; Absiella:1; Ruminiclostridium_D:1; Clostridium_X:1; Romboutsia:1; Clostridium_N:1; Clostridium_A:1; Clostridium_AC:1; Massilioclostridium:1; Clostridium_AG:1; Acetivibrio:1; Clostridium_D:1; Clostridium_AH:1; Clostridium_K:1; Clostridium_AN:1; Clostridium_C:1; Ruminiclostridium_B:1; Clostridium_V:1; Clostridium_AK:1; Clostridium_AF:1; Ruminiclostridium_F:1; Clostridium_T:1; Ruminiclostridium_C:1; Clostridium_U:1 |
| *Mycoplasma* | 81 | 2.2 | Mycoplasmopsis_A:17; Mycoplasmopsis:15; Metamycoplasma:13; Mesomycoplasma:10; Mycoplasmoides:7; Eperythrozoon_A:7; Mycoplasmopsis_B:4; Eperythrozoon_B:3; Malacoplasma:2; Malacoplasma_A:1; Mycoplasma_J:1; Mycoplasma_H:1 |
| *Campylobacter* | 61 | 1.7 | Campylobacter_A:32; Campylobacter_D:20; Campylobacter_B:8; Corynebacterium:1 |
| *Paenibacillus* | 58 | 1.6 | Paenibacillus_B:8; Paenibacillus_C:7; Paenibacillus_G:6; Paenibacillus_A:6; Paenibacillus_E:4; Paenibacillus_J:3; Paenibacillus_Q:3; Paenibacillus_O:3; Paenibacillus_K:2; Paenibacillus_T:2; Paenibacillus_R:2; Paenibacillus_D:2; Paenibacillus_V:1; Paenibacillus_S:1; Paenibacillus_F:1; Paenibacillus_M:1; Paenibacillus_P:1; Paenibacillus_N:1; Paenibacillus_L:1; Paenibacillus_I:1; Paenibacillus_U:1; Paenibacillus_H:1 |
| *Helicobacter* | 38 | 1.0 | Helicobacter_E:8; Helicobacter_A:8; Helicobacter_C:7; Helicobacter_D:7; Helicobacter_B:3; Helicobacter_G:2; Helicobacter_F:2; Helicobacter_H:1 |
| *Enterococcus* | 34 | 0.9 | Enterococcus_B:12; Enterococcus_A:8; Enterococcus_C:4; Enterococcus_D:2; Enterococcus_E:2; Enterococcus_G:2; Enterococcus_F:1; Enterococcus_H:1; Enterococcus_I:1; Enterococcus_J:1 |
| *Desulfovibrio* | 32 | 0.9 | Desulfovibrio_F:6; Desulfovibrio_H:6; Pseudodesulfovibrio:4; Desulfovibrio_A:3; Desulfovibrio_B:2; Desulfocurvibacter:1; Desulfovibrio_L:1; Desulfovibrio_M:1; Desulfovibrio_O:1; Desulfovibrio_J:1; Desulfovibrio_C:1; Desulfovibrio_K:1; Desulfovibrio_P:1; Desulfohalovibrio:1; Desulfovibrio_N:1; Desulfovibrio_I:1 |

**Supp. Table 18**. Genomic parameters calculated across 24,706 species clusters and 145,904 genomes.

| | Species | | | | | Genomes | | | | |
|---|---|---|---|---|---|---|---|---|---|---|
| | *Median* | *Mean* | *Std. dev.* | *5th percentile* | *10th percentile* | *Median* | *Mean* | *Std. dev.* | *5th percentile* | *10th percentile* |
| **Genome size (Mb)** | 3.3 | 3.6 | 2 | 1 | 7.5 | 3.9 | 3.8 | 1.7 | 1.5 | 6.6 |
| **GC content (%)** | 50.6 | 51.1 | 12.7 | 31.2 | 70.6 | 50.3 | 48.5 | 11.8 | 32.1 | 67.4 |
| **No. CDSs** | 3076 | 3336.6 | 1751.8 | 983 | 6748 | 3701 | 3644.2 | 1537.6 | 1493.2 | 6100 |
| **Coding density (%)** | 88.9 | 88.7 | 3.4 | 82.9 | 93.7 | 87.7 | 87.6 | 2.9 | 83.2 | 92.4 |
| **No. contigs** | 60.5 | 118.5 | 157.1 | 1 | 458 | 75 | 124.6 | 147.6 | 1 | 431 |

**Supp. Table 19.** List of 87 genomes retained in genome dataset despite failing the QC criteria as they represent genomes of high nomenclatural or taxonomic significance (see Excel file).

**Supp. Table 20.** Quality score of genomes used for selecting species representatives.

| **Feature** | **Score adjustment** | **Rational** |
|---|---|---|
| Complete genome | +100 | Genomes comprised of a single contig for each chromosome and plasmid are likely to be largely free of assembly errors |
| CheckM completeness estimate | + completeness | More complete genome assembles are less likely to have undetected contamination and are more suitable for many downstream applications including circumscribing species clusters |
| CheckM contamination estimate | -5×contamination | Contaminated genomes are problematic for many downstream applications including circumscribing species clusters |
| NCBI type material | +200 | Genomes annotated as being assembled from type material, even for species that are only effectively published, are essential for nomenclatural and taxonomic consistency between genomic resources |
| NCBI representative | +10 | All else being equal, it is beneficial for selected representative genomes at GTDB and NCBI to be the same |
| Full length 16S rRNA gene | +10 | Many downstream applications benefit from a genome having a complete 16S rRNA gene |
| No. contigs | -5×(no. contigs)/100 | Assemblies comprised of an excessive number of contigs are more likely to have assembly errors and represent additional challenges for many downstream applications |
| No. ambiguous bases | -5×(no. ambiguous bases)/100000 | Assemblies comprised of an excessive number of ambiguous bases are more likely to have assembly errors and represent additional challenges for many downstream applications |
| Assembled from metagenome | -200 | Metagenome-assembled genomes are far more likely to contain contaminating DNA than isolate or single-amplified genomes |
| Assembled from single cell | -100 | Single-amplified genomes are more likely to have assembly artifacts than isolate genomes |
